# Neutralizing antibodies elicited by Nipah virus G-head nanoparticle target diverse sites and inhibit receptor binding and fusion

**DOI:** 10.1101/2025.02.16.638571

**Authors:** Dan Zhou, Yong Wang, Yanfeng Yao, Wenhua Kuang, Rao Cheng, Gan Zhang, Hang Liu, Xin Li, Sandra Chiu, Zengqin Deng, Haiyan Zhao

**Affiliations:** State Key Laboratory of Virology and Biosafety, College of Life Sciences, Wuhan University, Wuhan, Hubei, China; State Key Laboratory of Virology and Biosafety, Wuhan Institute of Virology, Chinese Academy of Sciences, Wuhan, Hubei, China; Center for Biosafety Mega-Science, Wuhan Institute of Virology, Chinese Academy of Sciences, Wuhan, Hubei, China; Department of Laboratory Medicine, The First Affiliated Hospital of USTC, Division of Life Sciences and Medicine, University of Science and Technology of China, Hefei, Anhui, 230026, China; Key Laboratory of Anhui Province for Emerging and Reemerging Infectious Diseases, Hefei, China; Hubei Jiangxia Laboratory, Wuhan, Hubei, China

**Author notes:** These authors contributed equally to this work (Dan Zhou, Yong Wang, Yanfeng Yao, Wenhua Kuang, Rao Cheng). Corresponding author.; (H.Z.).

**Keywords:** Henipavirus, neutralizing antibodies, attachment G protein, neutralizing mechanisms, antigenic determinants

## Abstract

Nipah virus (NiV) and Hendra virus (HeV), two highly pathogenic Henipaviruses (HNVs), pose a significant public health threat. The attachment glycoprotein (G) plays a crucial role in viral attachment and entry, making it an attractive target for vaccine and therapeutic antibody development. However, the antigenic landscape and neutralization sensitivity of the G protein remain poorly defined. Here, we systematically characterize 27 monoclonal antibodies (mAbs) elicited by NiV G head (G^H^) nanoparticle-immunized mice. Among these, 25 mAbs exhibit neutralizing activity against two major NiV strains, NiV-Malaysia and NiV-Bangladesh, with five mAbs also cross-inhibiting HeV infection. Although all mAbs target G^H^, they inhibit viral infection through distinct mechanisms, either by blocking receptor ephrin-B2 engagement with the G protein or by preventing viral membrane fusion. Notably, mAbs from two distinct groups conferred complete protection to hamsters against lethal NiV-Malaysia infection. Structural analysis of NiV G^H^ in complex with four representative Fabs reveals four non-overlapping epitopes, including two novel antigenic sites and one shared protective epitope identified across species. Our study provides a comprehensive map of the antigenic determinants of NiV G^H^, elucidates the relationship between epitopes, neutralization mechanisms, and protection, and offers new insights and opportunities for antibody-based therapies and vaccine development.

**Highlights:** Most mAbs induced in G head nanoparticle-immunized mice are potent neutralizers MAbs recognizing the G head block receptor binding or viral fusion

Structural analysis shows mAbs target four distinct epitopes on G head protein

Receptor competition site on G head is a shared protective antigenic site in human and multiple animals

## Introduction

Nipah virus (NiV) and Hendra virus (HeV) are two highly pathogenic zoonotic henipaviruses (HNVs) known to cause severe encephalitis and respiratory diseases in humans with a high case fatality rate ^1–3^. Since their first identification in the mid to late 1990s, recurrent NiV outbreaks have been reported in Bangladesh and India ^4,5^. Although HeV outbreaks have been sporadic in Australia, HeV spillover from bats to horses has been frequently reported, resulting in several human exposures ^6,7^. HNVs can be transmitted to humans through contact with infected animals (such as bats, pigs and horses) or humans, or consumption of contaminated food, underscoring the potential global health risk posed by HNVs ^8^. Although mortality rates of 50-100% from HNV infection have been documented, there are still no clinically approved vaccines or specific therapeutics available for humans against HNVs ^2,9^. Thus, there is an urgent need to develop prevention and therapeutic strategies against HNV infections.

HNVs are negative-sense, single-stranded paramyxoviruses that possess two viral membrane proteins: the attachment glycoprotein (G) and the fusion protein (F) ^10^. The G protein exists as a tetramer on the viral surface and is a type II membrane protein, which can be structurally divided into four regions: the N-terminal cytoplasmic tail, the transmembrane domain, the stalk, and the C-terminal head domain (G^H^) ^11^. The C-terminal head domains of NiV and HeV are responsible for viral attachment to target cells by interacting with the receptors ephrin-B2 or ephrin-B3 ^12–14^. The F protein is a trimeric type I fusion protein, composed of three domains: the C-terminal cytoplasmic tail, the transmembrane domain, and the N-terminal ectodomain, which can be further divided into a head domain (DI, DII and DIII) and a C-terminal stalk ^15–17^. It is believed that G and F proteins are associated with each other; however, the organization of these two proteins on the whole virion remains unclear. Previous studies suggest that receptor engagement with the G protein likely promotes a conformational transition in the G protein, subsequently triggering F protein-mediated viral fusion with cell membranes to initiate infection ^18–21^.

Since the G and F proteins play critical roles in viral entry into host cells, they serve as the primary antigens for humoral immune response and vaccine development. Both G- and F-based vaccines have been investigated against HNV infections ^7,9,22–24^, showing robust immune responses and protective activity in animal models challenged with HNVs ^7,9^. In terms of monoclonal antibody (mAb) development, both human and mouse anti-F mAbs have been characterized in vitro or in vivo, with neutralizing antibodies targeting diverse epitopes across all three domains of the F head protein ^16,25–29^. While several mAbs targeting the G protein have demonstrated significant in vitro neutralizing effects and in vivo protection against NiV infection ^30–32^, most anti-G mAbs inhibit HNV infections by targeting the receptor ephrin-B2 binding site. Therefore, our understanding of the epitopes landscape on the HNV G glycoproteins targeted by antibodies is incomplete, especially for those with different inhibitory mechanisms.

We recently designed a ferritin-based NiV G head nanoparticle (NiV G^H^-ferritin) and generated a panel of mAbs from mice immunized with NiV G^H^-ferritin nanoparticle ^24^. Here, we further investigate the epitopes and neutralizing mechanisms of these newly identified mAbs to characterize the antigenicity of the NiV G head protein. We found that most G-reactive mAbs exhibit potent neutralizing activity against two authentic NiV strains, representing the two major NiV lineages: NiV Malaysia (NiV_M_) and NiV Bangladesh (NiV_B_). Five mAbs from two distinct competition groups demonstrate cross-neutralizing activity against another highly pathogenic HNV, HeV. Mechanistically, the anti-NiV G mAbs inhibit viral infection through either direct or indirect receptor blockade or by fusion inhibition. Structural analysis of NiV G^H^ bound to representative Fabs from the four distinct groups reveals the molecular determinants of protection and defines one protective, public epitope recognized by four different species. Overall, our study highlights that NiV G^H^ is a promising antigenic target of neutralizing antibodies and that G^H^-based vaccination in nonhuman species can induce neutralizing antibody responses to epitopes shared with humans.

## Results

### Most anti-NiV G mAbs are potently neutralizers to NiV

We previously developed 27 NiV G-reactive mAbs from NiV_M_ G^H^ nanoparticle-immunized mice, of which 25 showed potent neutralizing activity against vesicular stomatitis virus (VSV)-based NiV Malaysia pseudovirus (VSV-NiV_M_) ^24^. Although these 27 mAbs were classified into four competition groups (groups 1-4), their binding properties and neutralization sensitivity against distinct henipaviruses (HNVs) remain unclear. To further characterize the mAbs, we first assessed their cross-reactivity against G head proteins from multiple henipaviruses using enzyme-linked immunosorbent assay (ELISA). Of the 27 mAbs, 26 displayed strong binding signals to the recombinant G head proteins (residues 176-602) from two representative NiV strains: NiV_M_ and NiV_B_, at concentrations of 0.5 μg/mL or 1 μg/m (**Figure S1A and S1B**). We also assessed the binding capacity of the mAbs to the ectodomain of the NiV_M_ G protein (residues 96-602, G^ecto^) and found that all the mAbs exhibited similar binding capacity to G^ecto^ as compared to the NiV_M_ G^H^ protein (**Figure S1C**). Additionally, six mAbs were able to recognize G^H^ from the heterologous HeV (**Figure S1D**). None of the mAbs cross-reacted with G^H^ from two other related henipaviruses, Langya virus (LayV) or Mòjiāng virus (MojV) (**Figure S1E and S1F**).

Next, we evaluated the neutralizing activity of the mAbs against NiV_B_ and HeV pseudoviruses, as well as authentic viruses (**Figure 1**; **Table 1**). Of the 27 mAbs, 25 effectively inhibited infection of the NiV_B_ pseudovirus, with half-maximal inhibitory concentration (IC_50_) values of less than 0.3 μg/mL. Most mAbs (23 out of 27) were particularly potent, with IC_50_ values ≤ 0.015 μg/mL (**Figure 1A**). For HeV pseudovirus neutralization, five mAs exhibited varying levels of neutralizing activity, with IC_50_ values ranging from 0.001 to 9.86 μg/mL (**Figure 1B and 1C**; **Table 1**). Consistent with the pseudovirus neutralization results, 25 of the 27 mAbs potently inhibited authentic NiV_M_ and NiV_B_, and five also neutralized authentic HeV (**Figure 1D-1F**). Additionally, three previously reported mAbs (HENV-26, HENV-32, and nAH1.3) ^11,30^, which target distinct epitopes on G^H^, were included in both binding and neutralization assays. We previously demonstrated that HENV-26 belongs to group 3, nAH1.3 to group 4, and none of our mAbs competed with HENV-32 for G^H^ binding ^24^. Compared to the three control mAbs, 15 of our mAbs exhibited lower IC_50_ values against authentic NiV_M_, and 9 mAbs showed comparable or lower IC_50_ values against authentic NiV_B_ (**Figure 1**; **Table 1**). These findings suggest that potently neutralizing mAbs are consistently elicited in mice immunized with NiV_M_ G^H^ nanoparticle.

**Figure 1.**
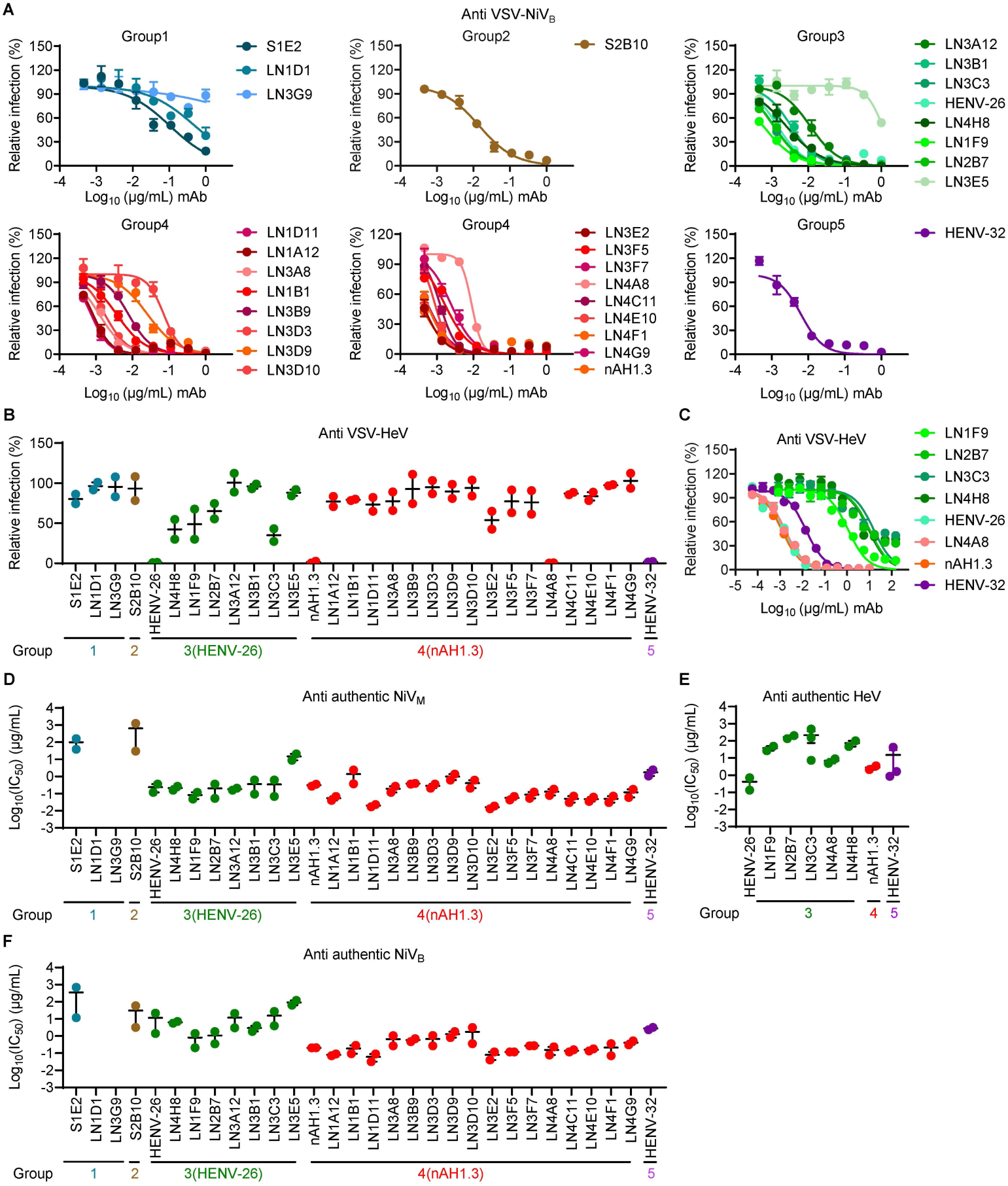
Neutralizing activity of NiV mAbs. (A) Neutralization curve plots of NiV mAbs against VSV-based NiV_B_ pseudovirus. (B) Neutralizing activity of NiV mAbs at 10 μg/mL against VSV-based HeV pseudovirus. (C) Neutralization curve plots of NiV mAbs against VSV-based HeV pseudovirus. (D-F) Neutralization activity of NiV mAbs against authentic NiV_M_ (D), HeV (E), or NiV_B_ viruses (F). Three independent experiments were performed in duplicate, and representative results are shown in (A) and (C). Data represented as mean ± SEM from two or three independent experiments performed in triplicate (B) or quadruplicate (D-F).

**Table 1.**
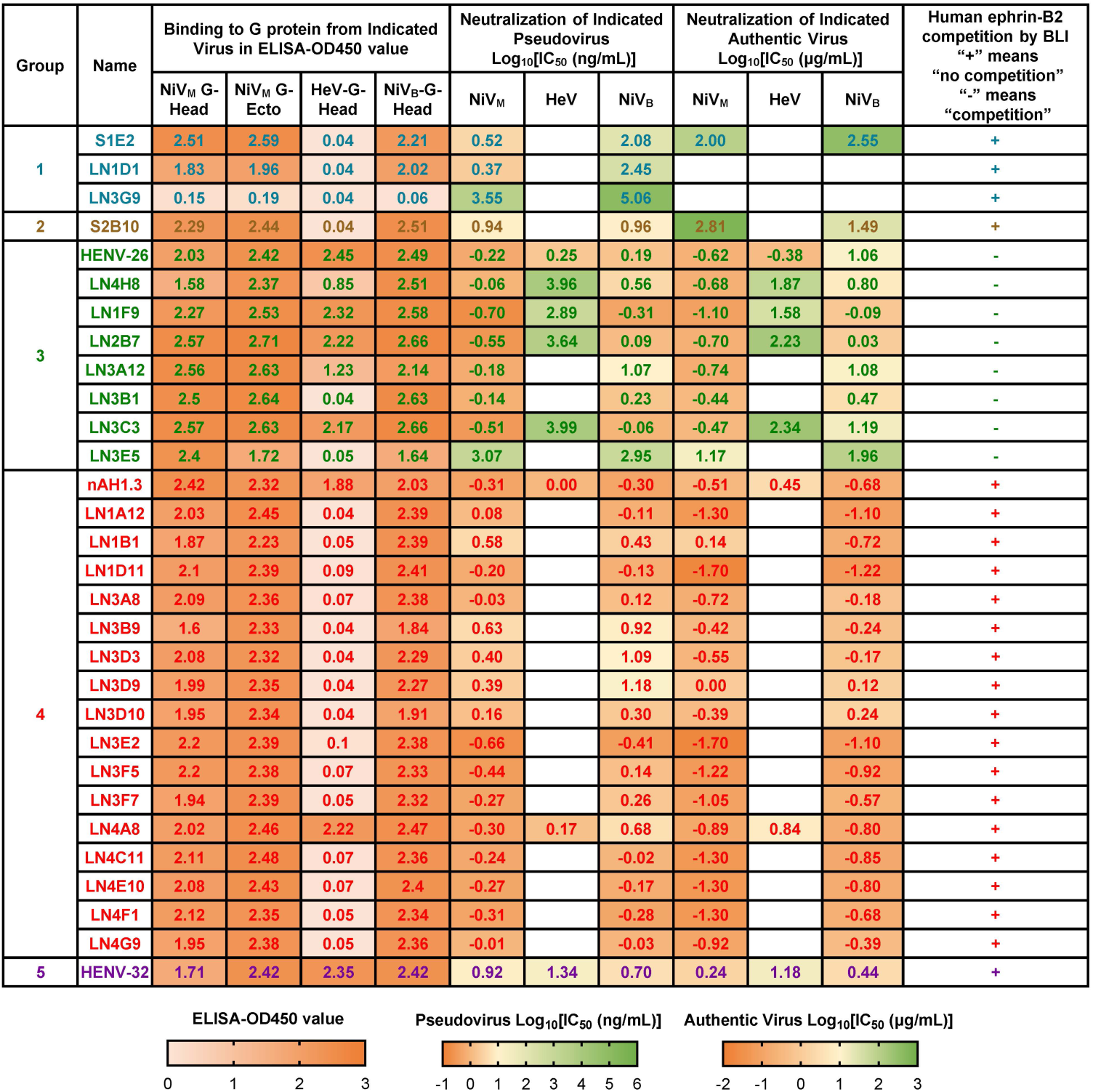
Binding and neutralizing activities of NiV mAbs. Two or three independent experiments were performed, and consistent results from all independent experiments were shown. IC50: 50% maximal inhibitory concentration.

### Neutralizing mAbs in groups 2, 3, and 4 block receptor binding via diverse mechanisms

Viral infection can be inhibited by mAbs at multiple stages, but it is particularly attractive and crucial for mAbs to interrupt viral infection at an early stage, before the virus begins to replicate. These include inhibiting receptor binding and viral-cell membrane fusion. To determine whether the mAbs block the host receptor from binding to the NiV attachment G protein, we performed a competition assay using Bio-Layer Interferometry (BLI) with recombinant NiV G^H^ protein. The BLI analysis showed that the binding of group 3 mAbs to G^H^ protein completely abolished the interaction between G^H^ and the ephrin-B2 receptor (**Figure 2A**). By contrast, ephrin-B2 was still able to efficiently bind to G^H^ in the presence of group 1, 2, or 4 mAbs. These results suggest that the group 3 mAbs inhibit virus infection through direct receptor binding inhibition, whereas the mAbs in the other three groups seem to neutralize NiV through distinct mechanisms.

**Figure 2.**
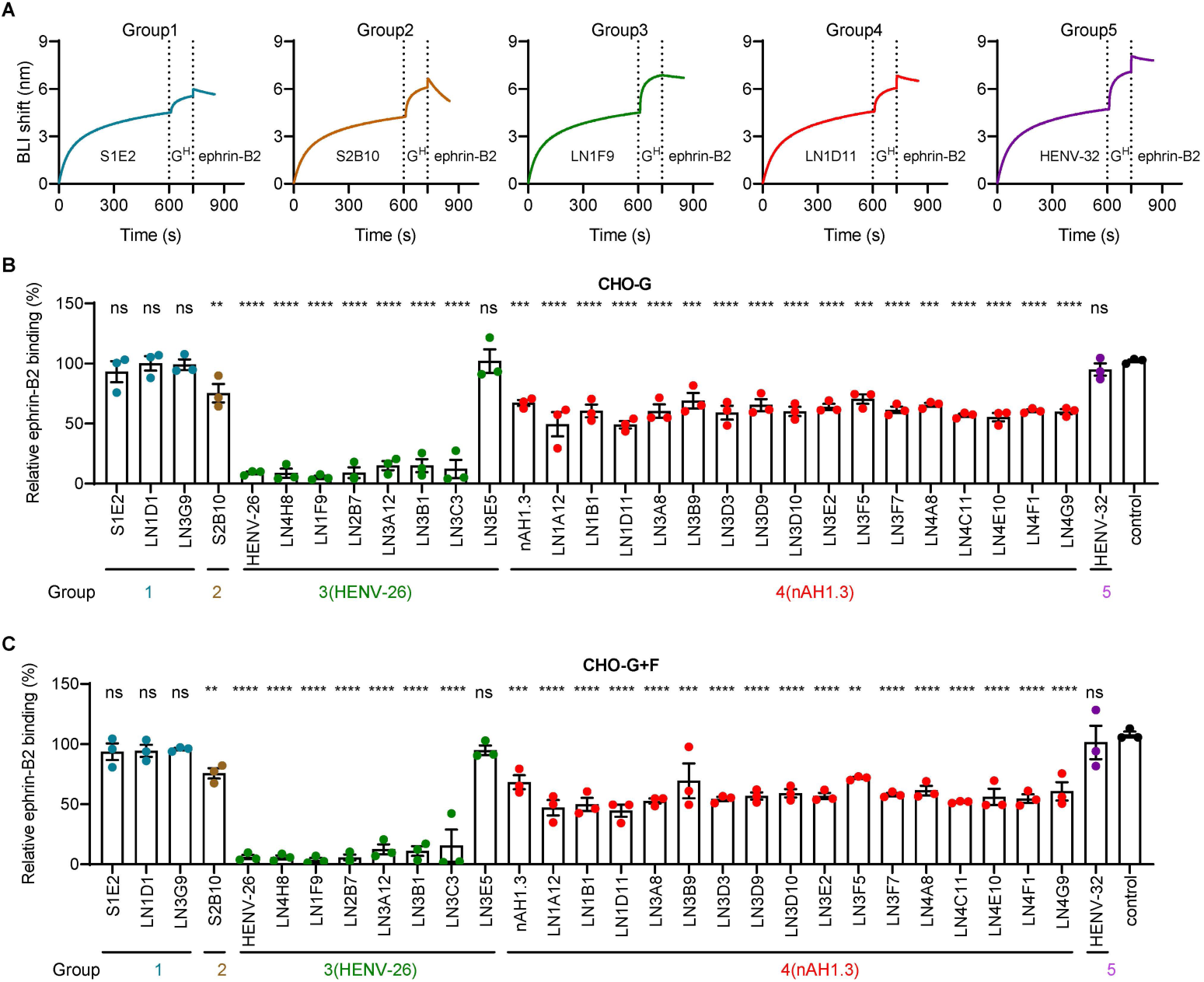
Competition between mAbs and the receptor ephrin-B2 for binding to NiV G. (A) The blocking activity of mAbs against host receptor ephrin-B2 binding to recombinant NiV G head protein was assessed by BLI. In this experiment, the indicated mAb was immobilized onto the biosensor to capture the NiV G head protein (G^H^) first, then the binding of the host receptor ephrin-B2 to captured G^H^ was measured. Parallel tips without antibody loading served as background controls. Representative curves (background subtracted) from two independent experiments are shown. (B-C) Competition flow cytometric assay showing binding of biotinylated ephrin-B2 to the full-length NiV G- or NiV G/F-expressing CHO cells in the presence of tested mAbs. The binding signal of ephrin-B2 to the G- or G/F-expressed cells without NiV mAbs treatment was set as 100%, and isotype mAb T3D9 was used as control. Data from three independent experiments performed in duplicate are presented as mean ± SEM. Statistical significance was determined by the one-way ANOVA with Dunnett’s multiple comparisons test, compared with control (T3D9) group. ns, not significant, \*\**p* < 0.01, \*\*\**p* < 0.001, \*\*\*\**p* < 0.0001.

Notably, our most potent neutralizing mAbs, including all five HeV cross-reactive mAbs, were derived from groups 3 and 4, which consist of 7 and 17 mAbs, respectively (**Figure 1**; **Table 1**). By contrast, group 2 contains only one mAb (S2B10), which did not block ephrin-B2 binding to recombinant G^H^ and exhibited only moderate inhibitory activity against NiV_M_ and NiV_B_. Group 1 mAbs demonstrated moderate to weak neutralization and similarly did not block ephrin-B2 binding to recombinant G^H^, as assessed by competition BLI.

We further assessed receptor binding inhibition by flow cytometry using full-length NiV G and NiV G/F-transfected cells to mimic the organization of NiV surface glycoproteins. Live NiV G or NiV G/F-expressing CHO cells were first incubated with individual mAbs and then stained with a fluorescence-labeled secondary antibody. Flow cytometric analysis showed that all 27 mAbs recognized G proteins displayed on the cell surface to varying degrees (**Figure S2**). We then analyzed the binding of ephrin-B2 to NiV G or NiV G/F-expressing CHO cells in the presence or absence of the mAbs (**Figure 2B and 2C**). Consistent with the BLI competition assay, pre-incubation of the cells with potent neutralizing mAbs in group 3 eliminated ephrin-B2 receptor binding, except for one modestly neutralizing mAb, LN3E5 (IC_50_=14.8 μg/mL against authentic NiV_M_). Unexpectedly, all group 4 mAbs (16 mAbs) decreased ephrin-B2 binding to cells by approximately 30%-50%, and the group 2 mAb S2B10 also showed a slight reduction in ephrin-B2 binding. However, no receptor binding blockade was observed for group 1 and group 5 mAbs. Together with the BLI competition results, we conclude that although neutralizing mAbs from groups 2 and 4 cannot directly block the binding of ephrin-B2 to recombinant G^H^ protein, their interactions with full-length G on the cell surface likely causes steric interference with ephrin-B2 binding.

### Neutralizing mAbs in group 1 and 5 inhibit viral fusion

To investigate whether the mAbs could inhibit viral fusion, we conducted a NiV glycoprotein-mediated cell-cell fusion assay using a dual split protein (DSP) system by monitoring renilla luciferase (Luc) activity (**Figure 3A-3E**) and green fluorescent protein (GFP) signal (**Figure 3F and 3G**). While mAbs from group 1 did not inhibit receptor binding (**Figure 2**), the neutralizing mAbs from this group (S1E2 and LN1D1) effectively reduced syncytium formation and cell-cell fusion with lowest 50% effective concentration (EC_50_) value of 287 ng/mL (**Figure 3A**). Despite group 2 mAb (S2B10) likely had steric interference with receptor binding to the G protein similarly as mAbs from group 4, S2B10 did not inhibit NiV glycoprotein-mediated cell-cell fusion (**Figure 3B**). By contrast, mAbs from group 4 exhibited potent fusion blockade activity similar to S1E2 from group 1 (**Figure 3A and 3D**). Although group 3 mAbs strongly inhibited receptor binding to the G protein, they only demonstrated weak fusion inhibition activity (~30%-50% decreases fusion) at high concentrations (1-10 μg/mL) (**Figure 3C**). HENV-32 (group 5) also inhibited fusion potently (**Figure 3E**), but did not show any blockage activity for ephrin-B2 binding (**Figure 2**). Consistent with the Luc activity, 1 and 10 μg/mL of mAbs exhibited similar fusion inhibition profiles as measured by GFP intensity (**Figure 3F and 3G**).

**Figure 3.**
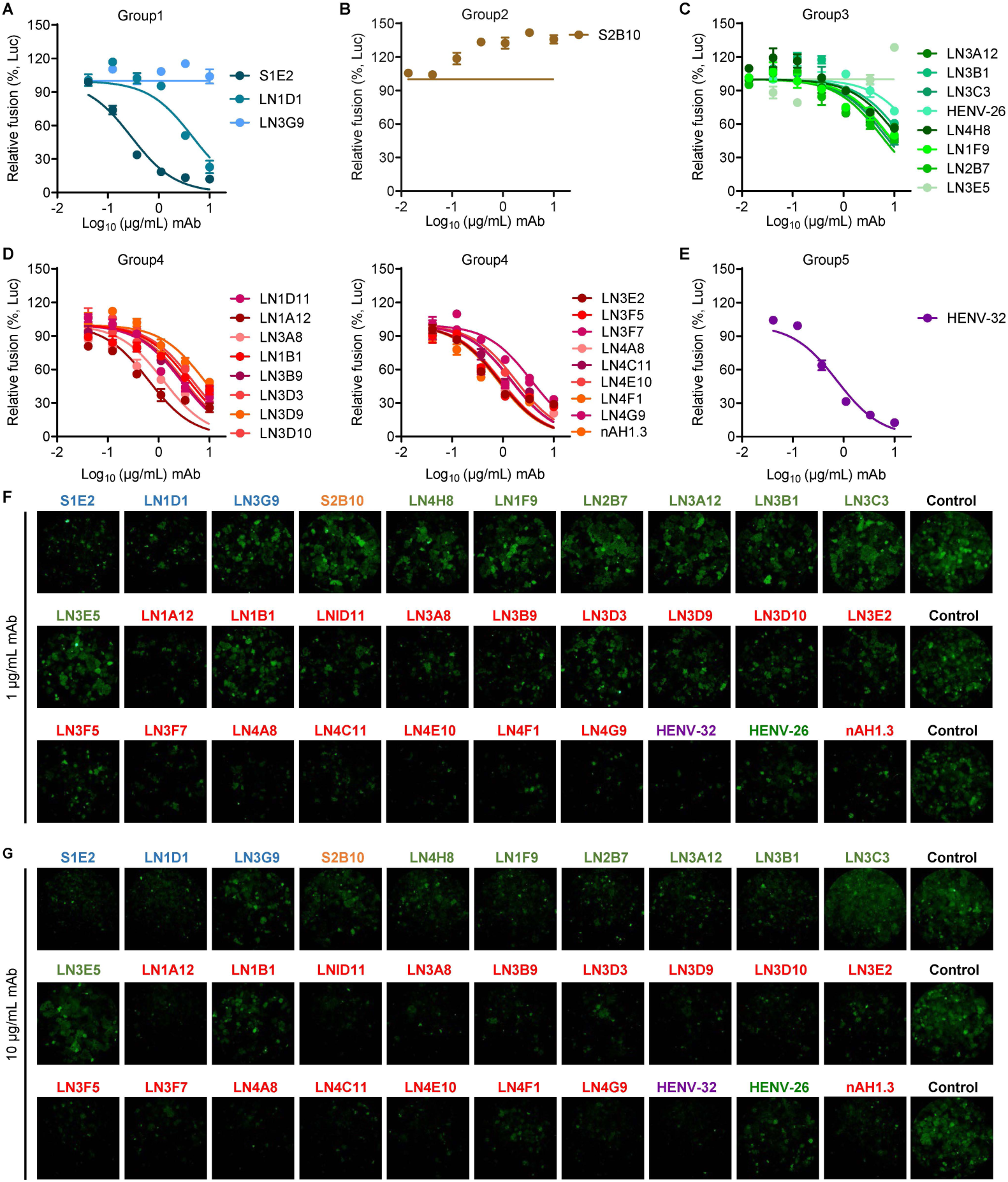
The fusion inhibition activity of NiV mAbs. NiV G/F-mediated cell-cell fusion was determined by a dual split protein (DSP) assay in 293T cells. (A-E) Fusion inhibition profiles of NiV mAbs quantified by luciferase activity. Three independent experiments were performed in duplicate, and representative results are shown. Each symbol represents the mean ± SEM. (F-G) Fusion activity in the presence of NiV mAbs from groups 1-5 at 1 μg/mL (F) or 10 μg/mL (G) was quantified by GFP signal. Three independent experiments were performed in duplicate, and representative results are shown.

### Potently neutralizing mAbs display in vivo protection against lethal NiV infection

We next assessed the protective activity of representative mAbs from each group against lethal NiV_M_ infection in vivo, both as pre-exposure prophylaxis and post-exposure treatment (**Figure 4**). For the pre-exposure prophylaxis, six-week-old Syrian golden hamsters were administered 30 mg/kg of antibody of either the tested mAb or an isotype control mAb S2A5 (an SFTSV mAb ^33^) intraperitoneally (i.p.) 24 hours prior to NiV infection^33^. The survival rate and body weight of the hamsters were subsequently monitored. All animals that received LN1F9 (group 3 mAb) or LN1D11 (group 4 mAb) survived without observed weight loss over the 14-day observation period. By contrast, all hamsters in the isotype control-treated group succumbed to the infection within 6 days post-infection (dpi). The administration of mAb S1E2 (group 1 mAb) or S2B10 (group 2 mAb) conferred partial protection against NiV challenge, as evidenced by the extended survival time of the hamsters and the fact that 1 out of 5 mAb-administered hamsters remained alive at the end of the study (**Figure 4B and 4C**).

**Figure 4.**
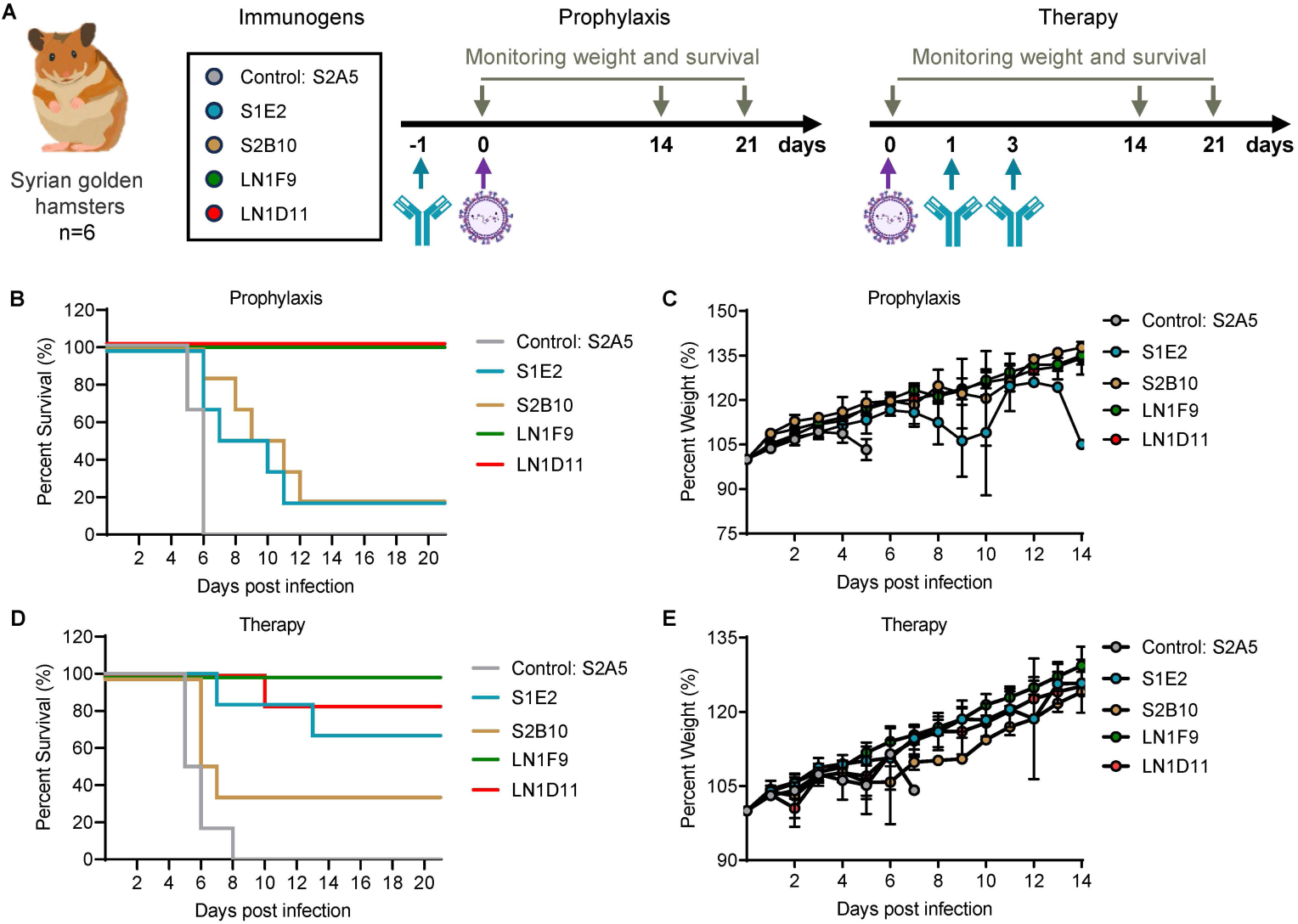
Anti-G mAbs display protection in hamsters against NiV infection. (A) Prophylactic or therapeutic treatment schemes of NiV mAbs in hamsters. For prophylactic treatment, hamsters (n=6) were administered the indicated mAbs at 30 mg/kg via the intraperitoneal route and then challenged with the NiV_M_ strains at 1000 LD_50_ via an intraperitoneal route 24 h later. For therapeutic treatment, hamsters (n=6) were challenged with the NiV_M_ strains at 1000 LD_50_ via an intraperitoneal route and then administered the indicated mAbs via the intraperitoneal route at 15 mg/kg twice (day 1 and 3 post-challenge). (B-C) Survival (B) and weight change (C) of Syrian hamsters during the prophylactic treatment regime. (D-E) Survival (D) and weight change (E) of Syrian hamsters during the therapeutic treatment regime. Data in each group are presented as mean and standard deviation (SD) in (C) and (E).

For post-exposure therapy, six-week-old hamsters were challenged with NiV, followed by the administration of the tested mAbs at 24 and 72 hours post-infection (a total of 30 mg/kg) via the intraperitoneal route (i.p.). Treatment with the non-binding isotype control mAb failed to protect the hamsters, with all animals succumbing to infection by 6-8 dpi. We observed 83%, 67%, and 33% survival rates in hamsters treated with LN1D11 (group 4 mAb), S1E2 (group 1 mAb), and S2B10 (group 2 mAb), respectively. However, only the group 3 mAb, LN1F9, exhibited 100% protection when administered as post-exposure therapy (**Figure 4D**). For the surviving hamsters, steady weight gain was observed across all mAb treatment groups throughout the course of the experiment (**Figure 4E**).

### Anti-NiV G mAbs recognize four distinct antigenic sites on G^H^

To gain insights into G-specific mAbs neutralization, we selected a representative mAb from each group (S1E2 from group 1, S2B10 from group 2, LN1F9 from group 3, and LN3D3 from group 4) and determined the complex structures of the G head protein (G^H^) bound to the antigen-binding fragments (Fabs) of these mAbs using cryo-electron microscopy (cryo-EM) or X-ray crystallography. Initially, we aimed to determine the structure of the G^H^ protein complexed with all four Fabs simultaneously, as epitope binning experiments suggest that these Fabs could bind concurrently. However, obtaining a high-resolution quinary complex structure proved challenging due to the preferred orientation of the sample. Consequently, we determined the cryo-EM structure of G^H^ bound to three Fabs (S1E2, S2B10, and LN3D3) at a resolution of 3.0 Å (**Figure S3 and Table S1**), and the crystal structure of G^H^ complexed with the LN1F9 Fab at 3.0 Å resolution (**Table S2**).

The quaternary complex structure reveals that the three Fabs could bind to G^H^ simultaneously without steric clash (**Figure 5A**), consistent with the competition biolayer interferometry results ^24^. N-linked oligosaccharides were observed on the G^H^ protein in the cryo-EM map, but none are located within the G-Fab interfaces. The surface areas on G^H^ buried by S1E2, S2B10, and LN3D3 are 764.3 Å^2^, 1007.7 Å^2^, and 900.9 Å^2^, respectively (**Figure 5C**).

**Figure 5.**
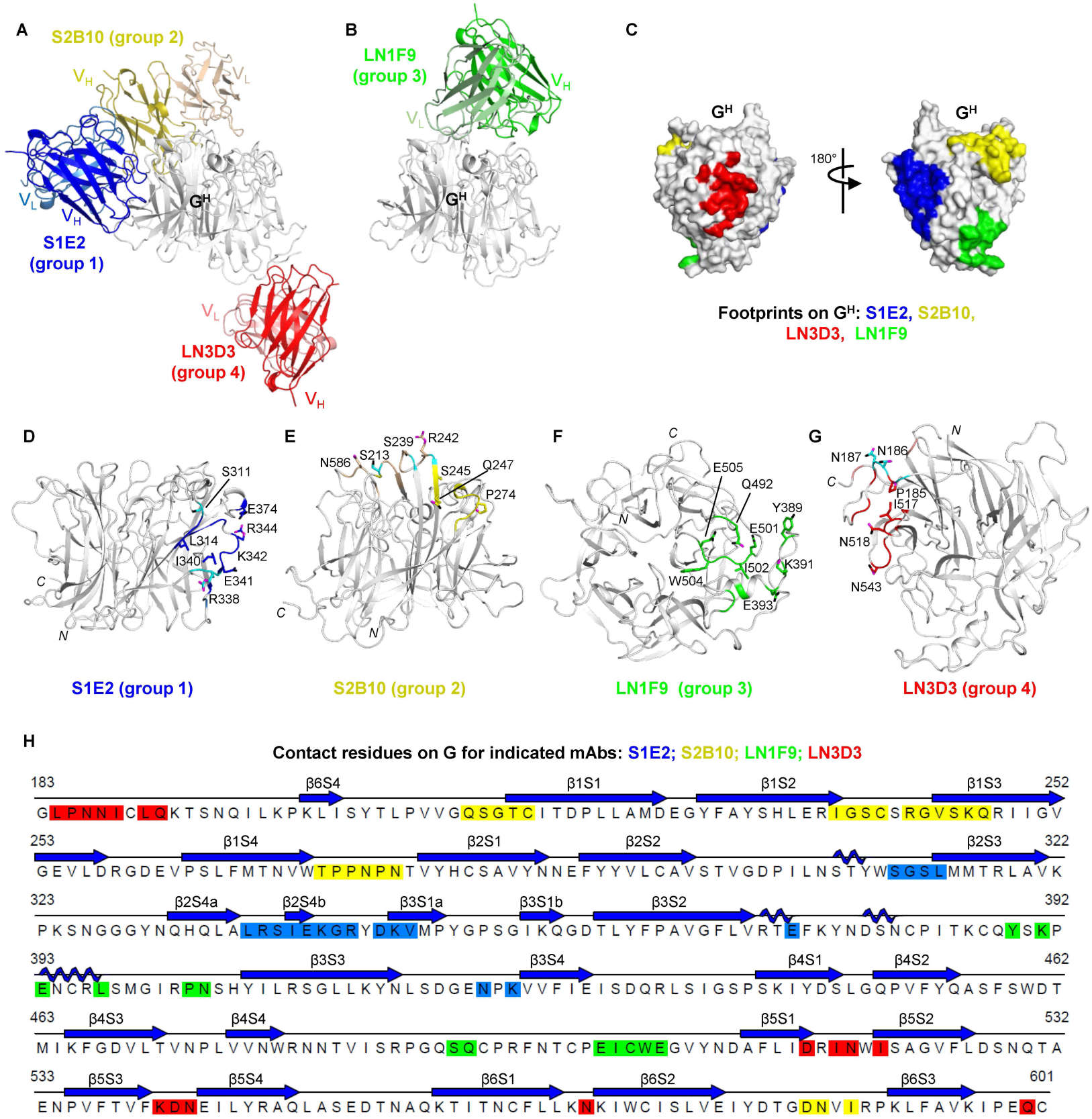
Structures of four representative Fabs in complex with NiV G^H^. (A) Cryo-EM structure of G^H^ bound simultaneously with Fabs of S1E2, S2B10 and LN3D3 is shown as a ribbon diagram. (B) Crystal structure of LN1F9 Fab complexed with G^H^ is shown as a ribbon diagram. The heavy and light chain variable domains of S1E2, S2B10, LN3D3 and LN1F9 are colored blue and skyblue, olive and wheat, red and salmon, green and palegreen, respectively. G^H^ is colored gray. (C) Surface representation of G^H^ with contact surfaces colored as in (A-B). (D-G) Delineation of the epitope contacts on the G^H^ for S1E2 (D), S2B10 (E), LN1F9 (F) and LN3D3 (G). Side chains are shown as sticks and labeled if they form hydrogen bonds, with nitrogen and oxygen atoms of the side chains colored magenta and black, respectively. G^H^ residues contacted by both chains are colored cyan. (H) The NiV_M_ G^H^ amino acid sequence with structurally defined epitopes highlighted as in panels (A-G).

Specifically, S1E2 interacts with the side of the G protein’s β-propeller (**Figure 5D and 5H**), engaging 18 residues distributed across four discontinuous segments: S311-L314, 11 residues on β3S1a strand and connecting loop of β2S4a-β3S1a (L337-R344 and D346-V348), E374, and two residues in the β3S3-β3S4 loop (N423 and K425). To corroborate the structural findings, we generated site-directed alanine substitution of selected residues in NiV G and assessed the binding ability of S1E2 to full-length G-expressing cells by flow cytometry. S1E2 showed markedly decreased binding when alanine mutation was introduced at epitope residue K342 (**Figure S4A**). Analysis of S1E2 contact residues on G protein reveals that the interaction is predominantly mediated by heavy chain, which contributes ~94% contacts (17 out of 18 residues), including four residues (S311, L337-S339) contacted by both the heavy and light chains (**Table S3 and S4**).

S2B10 binds to the top of the G protein β-propeller by engaging 24 residues across four discontinuous elements: the N-terminal region (Q212-C216), 10 residues in the β1S2-β1S3 loop and the N-terminal of the β1S3 strand (I237-C240 and R242-Q247), the β1S4-β2S1 loop (T272-N277), and the β6S2-β6S3 loop (D585, N586 and I588) (**Figure 5E and 5H**). Consistent with the structural analysis, S2B10 exhibited reduced binding phenotypes (~65%) to variant V244A (**Figure S4B**). Both the heavy and light chains contribute almost equally to the interactions, with 10 residues engaged by the heavy chain, 11 residues by the light chain, and 3 residues by both heavy and light chains (**Table S3 and S4**).

LN3D3 contacts the bottom loops of the G protein, including the N-terminal linker (L184-I188 and L190-Q191) as well as the C-terminal residue Q600, loop of β5S1-β5S2 (D515, I517-N518, and I520), loop of β5S3-β5S4 (D542-N543) and loop of β6S1-β6S2 (N570) (**Figure 5G and 5H**). Consistently, a single alanine substitution of N186 in G protein diminished >80% interaction of LN3D3 with G-expressing cells (**Figure S4D**). All six complementarity-determining regions (CDRs) from the heavy and light chains of LN3D3 are involved in G^H^ interaction, with the heavy chain contributing 60 % of the buried surface at the interface of the G^H^-LN3D3 complex (**Table S3 and S4**).

The crystal structure of G^H^-LN1F9 demonstrates that LN1F9 binds to the top loops of the G protein (**Figure 5B and 5F**). Docking of the G^H^-LN1F9 complex onto the quaternary cryo-EM structure of G^H^-S1E2-S2B10-LN3D3 indicates that the four mAbs target non-overlapping epitopes on G^H^, allowing all 4 mAbs to bind G^H^ simultaneously (**Figure 5C**). The epitope recognized by LN1F9 includes residues Y389, K391, E393, L397, and P403-N404, which are located within the β3S2-β3S3 loop, as well as residues S491-Q492 and E501-E505 in the β4S4-β5S1 loop (**Figure 5F and 5H**). Consistently, mutating E501 and E505 to alanine in G protein resulted in partial loss-of-binding phenotype, and LN1F9 further completely lost binding to a G protein variant with four amino acid substitutions: E391A/E393A/E501A/E505A (**Figure S4C**). Although LN1F9 interacts a total of 12 residues of G protein, all contacts are engaged by the heavy chain, with only one single residue contacted by both light and heavy chains. The surface area on G^H^ buried by LN1F9 is 626.6 Å^2^, with 90% of this area covered by the heavy chain (**Table S3 and S4**).

### Group 1 and 2 mAbs target two novel epitopes on NiV G^H^

To understand the antigenic features of G protein, we compared the epitopes recognized by our mAbs from four groups with the receptor ephrin-B2 and structurally available antibodies (**Figure 6**). Alignment of the structures of ephrin-B2-bound and mAb-bound NiV/HeV G with our G^H^-LN1F9 and G^H^-S1E2-S2B10-LN3D3 complexes revealed that G^H^-specific mAbs target at least six distinct antigenic regions on the NiV/HeV G protein. Notably, two of these epitopes, targeted by S1E2 (group 1) and S2B10 (group 2), represent novel sites that have not been previously described (**Figure 6**).

**Figure 6.**
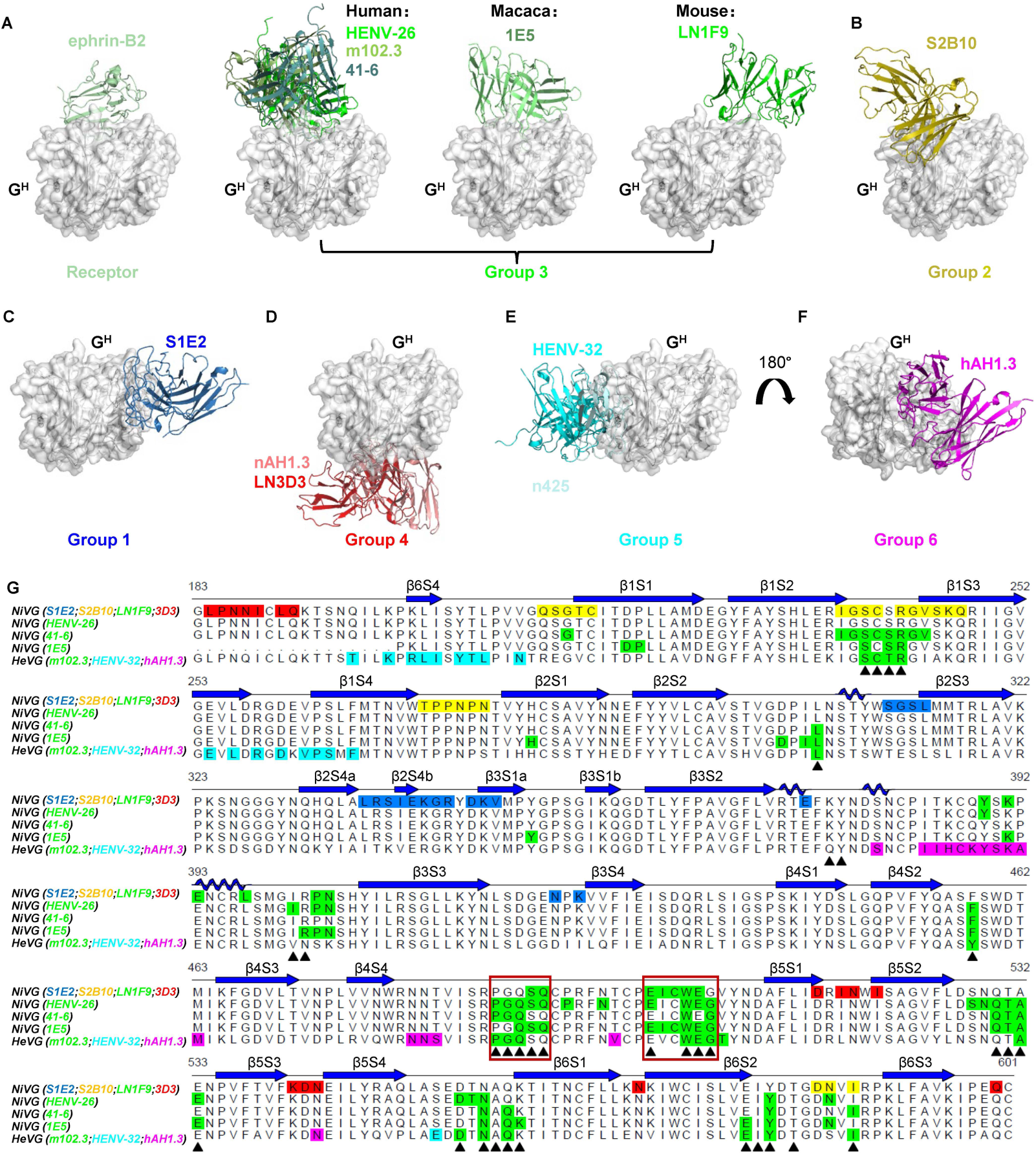
Group 3 mAb LN1F9 shares epitopes with human and macaca mAbs. (A-F) A comparison of structurally characterized mAbs targeting the G^H^ of NiV and HeV is shown. The mAb groups were defined based on the superimposition of the complexes, using G^H^ as a reference. Receptor ephrin-B2 (PDB: 2VSM) is displayed as a cartoon and colored palegreen. Group 3 mAbs: HENV26 (PDB: 6VY5), m102.3 (PDB: 6CMG), 41-6 (PDB: 8K3C), 1E5 (PDB: 8K0D) and LN1F9 are shown as cartoons in green, teal, lightteal, smudge and green, respectively. Group 1 mAb S1E2 is in blue, group 2 mAb S2B10 is in yellow, group 4 mAbs LN3D3 and nAH1.3 (PDB: 7TXZ) are in red and salmon, group 5 mAbs HENV32 (PDB: 6VY4) and n425 (PDB: 8XPS) are in cyan and palecyan, and group 6 mAb hAH1.3 is in magenta. (G) Sequence alignment of G^H^ from NiV and HeV, with structurally defined epitopes highlighted. The binding footprints of group 3 mAbs are colored green and the other groups are colored as in panel (A). The amino acids recognized by ephrin-B2 are marked with a black triangle below the sequence, and the common footprints recognized by ephrin-B2 and group 3 mAbs are indicated with red boxes.

The footprints of group 2 mAb (S2B10) are near the receptor ephrin-B2 binding site and also reside on the top of the G propeller (**Figure 6A and 6B**). Detailed analysis the binding footprints shows that several S2B10 contacts on G^H^ are also recognized by the ephrin-B2 and some group 3 mAbs (**Figure 6G**). In comparison to S2B10, the ephrin-B2 and group 3 mAbs bind to G^H^ from different orientations, and S2B10 binding to G protein results in varying degrees of steric clash with ephrin-B2 and group 3 mAbs (**Figure S5**). Group 1 (S1E2), group 5 (HENV-32 and n425), and group 6 (hAH1.3) target distinct regions on the side of the G propeller ^30,34,35^. Group 4 mAbs (LN3D3 and nAH1.3) target the bottom side of the G propeller, opposite to the ephrin-B2 binding site (**Figure 6C-6G**) ^11^. Interestingly, some mAbs from groups 3, 4, and 5 displayed cross-reactivity with the G proteins of both NiV and HeV, with potent cross-neutralization observed. In contrast, the epitopes recognized by mAbs from groups 1, 2, and 6 appeared to be specific to either NiV or HeV ^35^.

### Group 3 mAbs share epitopes with human and macaca mAbs

Group 3 mAbs target the top side of the G propeller, overlapping greatly with the footprint of receptor ephrin-B2, and inhibit HNV infection through mimicking receptor binding (**Figure 6**). This group includes one macaca-derived mAb (1E5) and three human mAbs (HENV-26, m102.3, and 41-6) ^30,31,36,37^. Structural analysis revealed that both the human and macaca mAbs utilize their relatively long CDR-H3 loops (18-23 amino acids) to insert into the central cavity of the G propeller, engaging the interior positions of the interface in a manner similar to ephrin-B2. By contrast, our mouse mAb LN1F9, with a shorter CDR-H3 loop (13 amino acids), only marginally protrudes into the G propeller’s cavity (**Figure S5**).

While all of the epitopes recognized by group 3 mAbs overlap significantly and directly block ephrin-B2 binding, each mAb also has unique interactions on the G^H^. Structural comparison further revealed two common loops (^488^PGQSQ^492^ and ^501^EICWEG^506^) for group 3 mAbs recognition of the ephrin-B2 binding site, suggesting that these two regions may represent a public epitope across species. These two loops form extensive hydrogen interactions with ephrin-B2 (5 hydrogen bonds) and group 3 mAbs (3-7 hydrogen bonds) ^31,35,37^, except for mAb 41-6 (one hydrogen bond) ^36^. Importantly, like other group 3 mAbs, LN1F9 demonstrated protective efficacy in NiV-challenged animal models and showed cross-neutralization against both HeV and NiV, highlighting the conservation and protective nature of this epitope across species.

### The receptor-binding site recognized by group 3 mAbs is an immunodominant antigenic site in mice and hamster

Syrian golden hamster is a well-studied animal model for HNVs infection and vaccine evaluation prior to the pre-clinal trial of anti-HNVs vaccines. We then assessed the antibody composition in sera from mice and hamsters immunized twice with NiV sG^H^ and NiV G^H^-ferritin by serum competition ELISA assay. 8 mAbs from five NiV G-bound groups (group 1: S1E2 and LN1D1; group 2: S2B10; group 3: LN3B1 and LN3A12; group 4: LN3E2 and LN3D3; group 5: HENV-32) were selected, and we monitored the binding signal of antisera from immunized animals to the NiV G head protein in the presence or absence of indicated mAbs (**Figure 7**).

**Figure 7.**
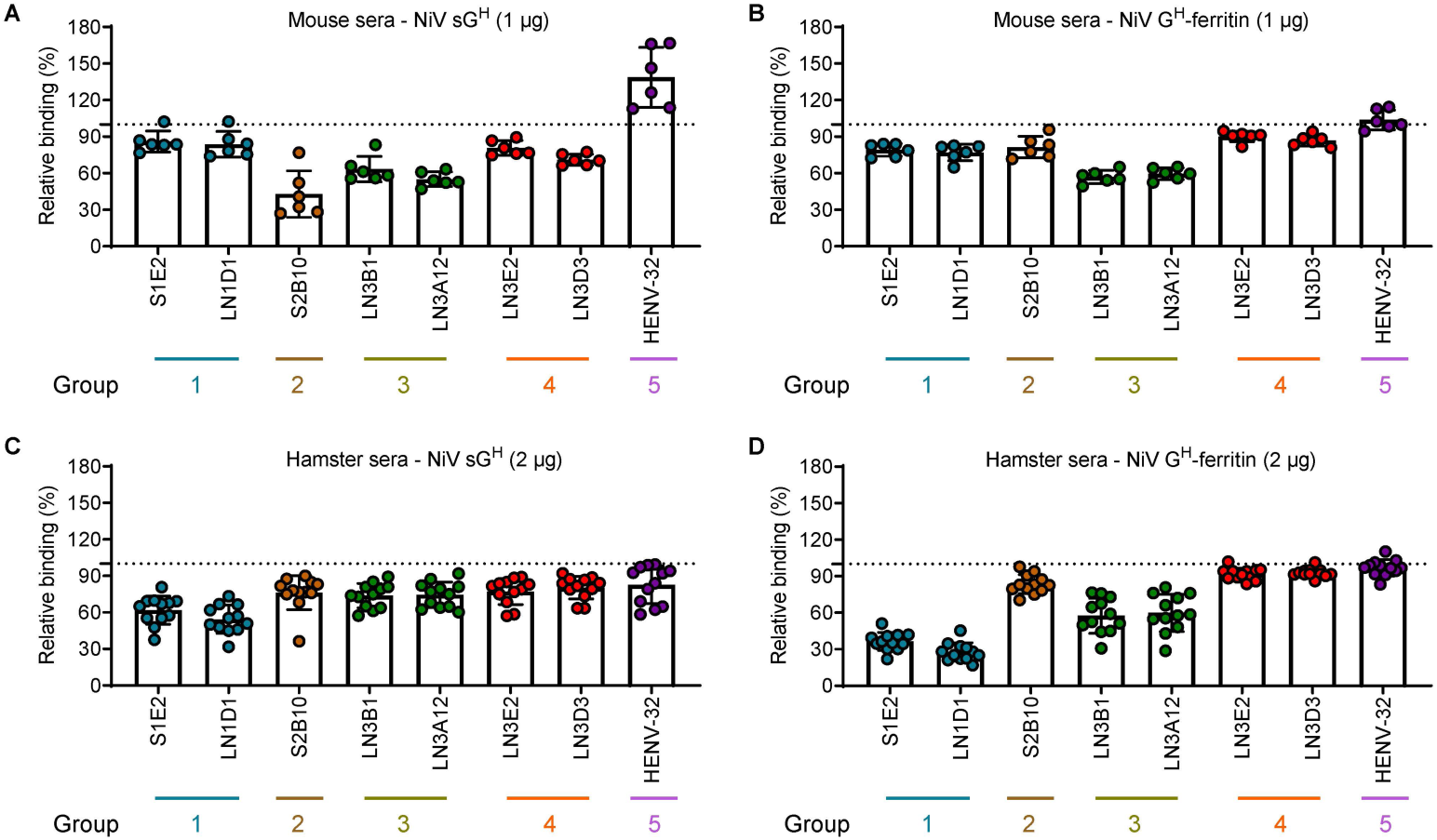
Quantification of the mAb proportion in sera from vaccinated mice and hamsters. (A-D) ELISA showing the binding of NiV sG^H^ (A and C) or NiV G^H^-ferritin (B and D) vaccinated sera to the immobilized NiV_M_ G head domain protein in the presence of 10 μg/mL of mAbs from group 1 (S1E2 and LN1D1), group 2 (S2B10), group 3 (LN3A12 and LN3D3), group 4 (LN3E2 and LN3D3) and group 5 (HENV-32). The binding of the sera in the presence of an isotype control mAb T3D9 (anti-SFTSV mAb, unpublished) was used as a control, and the OD450 (absorbance at 450 nm) value was defined as 100%. Each graph represents the mean and SD from two independent experiments performed in duplicate.

Serum competition analysis showed that NiV sG^H^ and NiV G^H^-ferritin could induce polyclonal antibodies of group 1-4, but not group 5 antibodies in both 1 μg/dose or 10 μg/dose immunized mice (**Figure 7A and 7B**). When preincubated G^H^ with mAbs first, the sera of NiV sG^H^ immunized mice lost ~15%, ~60%, ~40% and ~25% binding to G^H^ in the presence of mAbs from groups 1-4 (**Figure 7A**). For NiV G^H^-ferritin immunized mouse serum response, group 3 mAbs demonstrated the strongest binding inhibitory activity (**Figure 7B**). In hamster, NiV sG^H^-elicted sera recognize all five antigenic sites (**Figure 7C**), whereas NiV G^H^-ferritin-elicited sera primarily target antigenic sites within groups 1-3 (**Figure 7D**). This suggests that the antigenicity of the G head domain differs when presented as a nanoparticle compared to its soluble form. Additionally, group 1 and 3 mAbs showed dominant blockade activity against sera from NiV G^H^-ferritin immunized hamster. Combining these findings with mAbs isolated from mice, macaques, and humans, we conclude that group 3 mAbs are readily elicited in NiV G-immunized hamsters and mice, HeV G-vaccinated humans ^30^, and NiV/HeV G-immunized macaques ^31^, indicating that the epitope targeted by the receptor ephrin-B2 and group 3 mAbs represents an immunodominant ‘public epitope’ recognized across species.

## Discussions

Although several studies have generated mAbs targeting the G proteins of HNVs, these mAbs were derived from HeV-G vaccinated human ^30^, naïve human or human single-domain antibody libraries ^34,36^, or G-immunized macaque ^31^. There remains a need to systemically map the epitope landscape and neutralization sensitivity of the HNV G proteins. In this study, we characterize 27 mouse mAbs elicited by NiV G^H^-ferritin both structurally and functionally, revealing two novel antigenic sites on G^H^ and one protective, shared epitope across multiple species. Our newly isolated mAbs, along with previously reported mAbs, recognize at least five distinct epitopes on NiV G^H^ and inhibit NiV infection by either blocking receptor recognition through epitope competition or steric hindrance, or by inhibiting fusion. This work provides antigenic knowledge of NiV G^H^ and uncovers the relationships between epitopes, functions, and the mechanisms of action of anti-NiV G mAbs.

Most mAbs in our panel exhibited potent neutralizing activity against both pseudotyped and authentic NiV strains (NiV_M_ and NiV_B_), and five mAbs also cross-neutralized pseudotyped and authentic HeV in vitro. Additionally, representative neutralizing mAbs from 4 groups demonstrated partial to complete protection against lethal NiV_M_ challenge in vivo, with groups 3 (LN1F9) and 4 (LN1D11) mAbs showing 100% protection as prophylaxis. Notably, all five HeV cross-reactive mAbs belong to groups 3 and 4, though no cross-reactivity was observed against two other potential human henipaviruses, Langya virus and Mòjiāng virus, likely due to their relatively low sequence identity (<30%). While further studies are needed, we speculate that the cross-protection against HeV by mAbs from these two groups is desirable. These findings further underscore G^H^ as a promising antigenic target for broadly neutralizing antibodies and vaccine development.

As their names suggest, the attachment glycoprotein (G) primarily mediates viral attachment to host cells, while the fusion glycoprotein (F) initiates viral fusion with the host cell membrane during NiV infection. Although our mAbs bind to four distinct epitopes on the NiV G protein with varying neutralizing potency, potent inhibitory mAbs from different groups can block either viral fusion or receptor recognition. A previously reported group 5 mAb, HENV-32, provided post-exposure protection against NiV challenge in ferrets, though the mechanistic basis of its protection remains unclear ^30^. We here demonstrate that HENV-32 has minimal effect on recombinant ephrin-B2 binding to NiV G, whether in its soluble form or displayed on the cell surface, mimicking the glycoprotein configuration on the virion surface. Similar to group 1 neutralizing mAbs, HENV-32 efficiently reduced NiV glycoprotein-mediated cell-cell fusion without affecting receptor recognition.

Structural analysis revealed that S1E2 (group 1 mAb) and HENV-32 (group 5 mAb) bind to distinct regions on the side of the G propeller, with HENV-32 likely causing a steric clash with the stalk or G head domain of adjacent G proteins. By contrast, S1E2’s epitope on the G tetramer is fully exposed, and no steric clash occurs when it binds to any G subunit. Despite targeting different epitopes on the G protein, both mAbs share a common fusion inhibition mechanism, likely by preventing the conformational transition of G required to trigger fusion or by impacting the interaction between G and F. Similarly, many mAbs recognize distinct antigenic sites on viral structural proteins, yet exhibit similar attachment or fusion inhibition phenotypes, as seen in different viruses ^33,38–40^.

Group 5 mAbs have been reported in HeV G-vaccinated humans or naïve human single-domain library ^30,34^; however, we did not observe the presence of group 5 mAbs in the sera of NiV sG^H^- or G^H^ nanoparticle-immunized mice. Furthermore, none of the 27 mAbs showed competition with the group 5 mAb HENV-32 for NiV G^H^ binding, as detected by BLI, consistent with structural comparison data. Notably, 17 out of the 27 mouse mAbs belong to group 4, with only one mouse-derived mAb nAH1.3 from hybridomas previously reported falling into this group ^11,41^. While a larger panel of human and mouse mAbs is needed for further investigation, these findings suggest that mice do not fully recapitulate the human antibody response to HNV G proteins, likely due to differences in B cell repertoires between the two species.

The group 3 mAb LN1F9 provided both pre- and post-exposure protection against lethal NiV_M_ infection in hamsters. The group 4 mAb LN1D11 conferred complete protection when administered prophylactically and 83% protection when used therapeutically. Our structural and mechanistic studies reveal that mAbs in groups 3 and 4 bound non-overlapping epitopes on G^H^ and inhibited NiV infection through distinct strategies. Notably, reported mAbs from these two groups also exhibited cross-reactive and inhibitory activity against three pathogenic HNVs: NiV_M_, NiV_B_, and HeV. Since the amino acids on G^H^ recognized by three human mAbs significantly overlap with the epitope targeted by group 3 mAb LN1F9, this suggests that the epitope recognized by group 3 is a hotspot in humans, making it susceptible to escape mutants under group 3 antibody-imposed selection pressure. By contrast, the epitope targeted by group 4 mAbs appears to be mouse-specific and may experience relatively weaker selection pressure compared to the more dominant human epitopes. Therefore, a combination of mAbs from these two groups would be an effective countermeasure against HNV infections in the future.

In summary, our study reveals the antibody components and mechanisms of the humoral immune response at both monoclonal and polyclonal levels induced by NiV G^H^ immunogen, further demonstrating that NiV G^H^-ferritin can serve as a promising vaccine candidate against NiV infection. Our characterization of neutralizing mAbs, which exhibit distinct neutralization profiles against different HNVs, target diverse epitopes on G^H^, inhibit NiV infection at multiple stages, and demonstrate therapeutic potential, provides valuable insights for the rational design of vaccines against highly lethal HNVs.

## Limitations of the study

Although we demonstrate that G-specific mAbs from distinct groups could disrupt viral fusion, the detailed inhibitory mechanisms may vary. The absence of a G-F complex structure and a comprehensive understanding of the dynamic rearrangements of G required to trigger the transition of F from the pre-fusion to post-fusion stage hampers our understanding of the entry mechanisms and infection processes of HNVs. Our delineation of the antigenicity of the NiV G protein reveals several mAbs with in vivo protective activity against NiV infection; however, humanization of these mAbs with therapeutic potential is needed to improve their safety profile.

## Materials and Methods

### Ethics statement

Hamster experiments were approved by the Wuhan Institute of Virology, Chinese Academy of Sciences (approval number: WIVA21202301). Authentic virus infections were performed in the animal biosafety level 4 (ABSL-4) facility at the National Biosafety Laboratory (Wuhan), Chinese Academy of Sciences.

### Viruses and cells

The NiV_M_, NiV_B_, and HeV strains were obtained from the National Virus Resource Center at the Wuhan Institute of Virology for authentic virus neutralization assays and challenge studies. All authentic viruses were propagated in Vero E6 cells. Vero E6 and 293T cells were cultured at 37°C with 5% CO_2_ in DMEM (Monad Biotech) containing 8% or 10% FBS (ExCell Bio). CHO-K1 cells were cultured in DMEM/F12 (1:1) medium (Biosharp, Cat#: BL305A) with 10% FBS. Expi293 cells were cultured at 37°C with 8% CO_2_ in SMM 293-TII Expression Medium (Sino Biological, Cat#: M293TII) or 293F Hi-exp Medium (OPM Biosciences, Cat#: AC601501).

### Plasmids

DNA segments encoding the full-length NiV_M_ G (GenBank: NP_112027.1), NiV_M_ F (GenBank: NP_112026.1), NiV_B_ G (GenBank: AAY43916.1), NiV_B_ F (GenBank: AAY43915.1), HeV G (GenBank: NP_047112.2), HeV F (GenBank: NP_047111.2), MojV G (GenBank: YP_009094095.1), LayV G (GenBank: UUV47206.1) and human ephrin-B2 (residues 27-167; GenBank: NP_004084.1) were codon-optimized for expression and synthesized by Tsingke Biotechnology Co. The coding regions for NiV_M_ G head domain (residues 176-602), NiV_M_ G ectodomain (residues 96-602), NiV_B_ G head domain (residues 176-602), HeV G head domain (residues 176-605), LayV G head domain (residues 176-625), and MojV G head domain (residues 176-625) were individually cloned into a mammalian expression vector with a signal peptide, an N-terminal 6×His-tag, and an N-terminal HRV 3C protease cleavage site. The coding regions for human ephrin-B2 (residues 27-167) were individually cloned into a mammalian expression vector with a signal peptide and a C-terminal 6×His-tag. Coding regions for the full-length G and F of NiV_M_, NiV_B_, and HeV were cloned into pCAGGS vector, respectively, for VSV-based pseudovirus generation. To increase the pseudovirus titer, S207L and G252D point mutations were introduced on the NiV_B_ F gene, and a truncated variant was introduced on the HeV F gene, leaving 5 residues in the cytoplasmic tail ^42^. The variable regions of heavy (VH) and light chains (VL) were cloned into AbVec2.0-IGHG1 (Addgene) and AbVec1.1-IgKC (Addgene) or AbVec2.1-IGLC2 (Addgene) expressing vectors, respectively, for mAb generation. To generate Fab, the VH segments were also cloned into a modified AbVec2.0-IGHG1 expressing vector with a C-terminal 6×His-tag after the CH1 constant region.

### Protein expression and purification

Proteins were generated in Expi293 cells by transient transfection using CarpTrans (OPM Biosciences, Cat#: AC501302). Briefly, 200 μg expression plasmids of human ephrin-B2, HNVs G, mAbs, or Fabs were mixed with 800 μL CarpTrans following the manufacturer’s protocol and added into 200 mL Expi293 cells. For Fabs and mAbs generation, the paired heavy- and light-chain plasmids were co-transfected into Expi293 cells at a molar ratio of 1:1 (for Fab) or 1:1.2 (for mAb). Five or six days after transfection, the cell supernatants containing proteins or Fabs were harvested and filtered through 0.45 μm filters. The Fabs were purified using Ni-Charged Resin (GenScript, Cat#: L00666) and size exclusion chromatography (Superose 6 Increase 10/300 GL column or Superdex 200 Increase 10/300 GL column). MAbs were purified using rProtein A Beads (Smart Lifesciences, Cat#: SA015100).

To generate the NiV_M_ G head protein for crystallization, 5 μM kifunensine (Toronto Research Chemicals, Cat#: K450000) was added to the culture medium before transfection. The purified protein was treated with Endo HF (New England Biolabs, Cat#: P0703L) and HRV 3C protease to produce a homogeneously deglycosylated NiV_M_ G head domain without the His-tag.

### Enzyme-linked immunosorbent assay (ELISA)

For binding activities of NiV G mAbs to different HNVs G proteins, 96-well ELISA plates (Corning, Cat#: 9018) were coated with 3 μg/mL of the purified HNV G head protein in coating buffer (0.1 M carbonate, pH 9.6) at 4°C overnight. The plates were then washed with PBS-T (PBS with 0.05% Tween 20) and blocked with blocking buffer (PBS-T containing 1% BSA) at 37°C for 2 h. After blocking, the buffer was replaced with fresh blocking buffer containing NiV G mAbs (0.5 μg/mL for NiV_M_ G head domain and NiV_M_ G ectodomain, and 1 μg/mL for other HNVs G head domain) and incubated at 37°C for 2 h. Following four washes, the plates were incubated with HRP-conjugated goat anti-human IgG (ABclonal, Cat#: AS002) at 37°C for 1 h. After four additional washes, the plates were developed with one-component TMB chromogen solution (NCM Biotech, Cat#: M30500) at 37°C for 10-30 min, and the reaction was stopped with 1 M HCl. The absorbance at 450 nm was measured using a Varioskan LUX (Thermo Scientific).

For the proportion of different group antibodies in sera from hamsters and mice, 3 μg/mL NiV_M_ G head domain was coated on the ELISA plates. After blocking, 10 μg/mL representative G-specific mAbs from groups 1-5 were used to occupy the corresponding epitope. Wells with 10 μg/mL T3D9 (an anti-SFTSV mAb, unpublished) served as controls, with their absorbance at 450 nm defined as 100%. After 90 min incubation at 37°C, 50 μL sera from NiV sG^H^ or NiV G^H^-ferritin vaccinated mice or hamsters, at appropriate dilution, were added into the ELISA plates and incubated at 37°C for 30 min. HRP-conjugated goat anti-hamster or anti-mouse IgG (Thermo Scientific, Cat#: PA128823; Cat#: 31430) were used for hamster or mouse sera detection, respectively.

### Biolayer interferometry assay (BLI)

The competition biolayer interferometry (BLI) assay was conducted on an Octet Red 96 device (Pall ForteBio) to evaluate whether NiV G mAbs can block the binding of human ephrin-B2 to purified NiV G^H^ protein. Briefly, 10 μg/mL of NiV G mAbs were first loaded onto ProA Biosensors (Sartorius, Cat#: 18-5010). After a 10-second wash with running buffer, the biosensor tips were dipped into wells containing 500 nM NiV_M_ G^H^ protein for 120 s. All proteins were diluted in running buffer (10 mM HEPES, 150 mM NaCl, 3 mM EDTA, 0.05% Tween-20, and 1% BSA, pH 7.4). Subsequently, the tips were then immersed in a buffer containing 10 or 100 μg/mL of human ephrin-B2 for 120 s to monitor the binding signal of ephrin-B2 to the mAb-captured G^H^. Tips without mAbs loading were run in parallel to define the background signal. Data were analyzed using Octet data analysis software (version 12.2.0.20).

### Pseudovirus packaging and neutralization assays

The VSV-based HNV (NiV_M_, NiV_B_, and HeV) pseudotyped viruses were generated following previously published protocols ^24,43^. Briefly, pCAGGS plasmids encoding the full-length HNV G and F genes were co-transfected into 293T cells using Gene Twin (Biomed, Cat#: TG101). After transfection, the cells were cultured for 16-24 h and then infected with VSVΔG-eGFP for 4-6 h. Following infection, the cells were washed with PBS and incubated with 1 μg/mL of anti-VSV G mAb I1 ^44^ diluted in DMEM with 4% FBS. 24-30 h post-infection, the pseudovirus supernatants were harvested and aliquoted before storage at −80°C.

For the pseudovirus neutralization assay, Vero E6 cells were seeded in the 96-well plates at a density of 1.5×10^5^ cells/mL the day before the experiments. Serially diluted NiV G mAbs or Fabs were mixed with pseudovirus and incubated at 37°C for 1 h. MAb or Fab-pseudovirus mixtures were then added to the Vero E6 cells. The following day, cells were fixed with 4% paraformaldehyde (PFA), and the green fluorescent dots were counted using a CTL-S6 Universal M2. IC₅₀ (half-maximal inhibitory concentration) was calculated using GraphPad Prism (v.8.0) with a nonlinear regression model.

### Flow cytometric assay

A flow cytometric assay was used to identify critical residues on NiV G for mAb binding. Expi293 cells were transiently transfected with wildtype NiV_M_ G (positive control), G mutant plasmids, or pCAGGS vector (negative control). At 48 hours post-transfection, the cells were fixed with 4% PFA for 10 min, and the cells were then washed and incubated with 10 μg/mL of the indicated NiV G mAbs at 4°C for 30 min. Following washing, the cells were stained with Alexa Fluor 488 anti-human IgG antibody (Thermo Scientific, Cat#: A-11013) at 4°C for 30 min. The cells were then washed again, resuspended, and subjected to analysis using a CytoFLEX S (Beckman). The binding capacity of the indicated NiV G mAbs to G point mutations was calculated relative to wildtype G and normalized by the G protein expression levels. Cells stained with 10 μg/mL of NiV G mAb-mix (mix of representative mAbs from 5 groups: S1E2, S2B10, LN1F9, LN3D3, and HENV-32, 2 μg/mL of each antibody) were used to calculate relative expression levels of G point mutations compared to the wildtype G.

For the competitive flow cytometric assay, CHO-K1 cells first were transfected with plasmids encoding full-length NiV_M_ G or co-transfected with plasmids encoding both full-length NiV_M_ G and F. At 24 hours post-transfection, the cells were dissociated from the plates using 5 mM EDTA. After washing, the cells were incubated with 10 μg/mL of the indicated NiV G mAbs, isotype control mAb T3D9 (an anti-SFTSV mAb, unpublished), or FACS buffer without mAb (negative control) for 30 min. Subsequently, 50 μg/mL of biotinylated human ephrin-B2 were added to the cells and incubated for 30 min, followed by washing and incubation with Streptavidin-APC (BD Biosciences, Cat#: 554067) for 30 min. The cells were then washed again, resuspended, and analyzed by flow cytometry (CytoFLEX S, Beckman). The binding of ephrin-B2 to the G- or G/F-expressing cells in the presence of mAbs was compared with negative control cells incubated with FACS buffer.

### Authentic virus neutralization assay

The NiV G mAbs were prepared in a three-fold serial dilution in DMEM containing 2% FBS and incubated with either 100 TCID_50_ NiV_M_, 100 TCID_50_ NiV_B_, or 100 TCID_50_ HeV for 1 h at 37°C. The virus-mAb mixtures were then added to Vero E6 cells in 96-well plates and incubated for 1 h at 37°C. After incubation, the cells were washed and cultured in DMEM supplemented with 2% FBS until the cytopathic effect was observed approximately five days post-infection. Each mAb dilution was set up in four replicates. IC_50_ values were calculated using IBM SPSS Statistics 27.

### Crystallization and structure determination of NiV G-LN1F9 Fab complex

The purified NiV_M_ G head domain was mixed with LN1F9 Fab in a molar ratio of 1:1.2, and the complex was further purified to homogeneity by size exclusion chromatography using a HiLoad 16/600 Superdex 200 pg column (Cytiva). Crystallization of the NiV G-LN1F9 complex was performed by sitting drop vapor diffusion at 16°C. Typically, 25 or 35 mg/mL protein was mixed with the precipitant/reservoir solution at a 1:1 volume ratio in a 0.6-μl drop. Crystals appeared in the precipitant/reservoir solution of 0.3 M ammonium formate, 0.1 M HEPES pH 7.0, and 20 % (vol/vol) Sokalan CP 5 within 1 week. Crystals were stepwise transferred to a cryostabilizer solution (precipitant solution supplemented with 35% [vol/vol] glycerol) and then flash-cooled in liquid nitrogen before data collection.

The X-ray diffraction data were collected at the BL10U2 beamline of Shanghai Synchrotron Radiation Facility (SSRF) with a wavelength of 0.9792 Å and temperature of 100 K. A total of 360 degrees of data were collected in 0.5° oscillation steps. The diffraction data were automatically processed by the pipeline Xia2 ^45^ at the beamline and scaled with Aimless ^46^ in the CCP4 suite ^47^. Phasing was obtained by molecular replacement using PHASER ^48^ with the crystal structure of NiV G^H^ (PDB ID: 7TXZ) and the AlphaFold2-predicted Fab model as search models ^49^.

### Cryo-EM sample preparation

NiV_M_ G head domain was incubated with a molar excess of S1E2, S2B10, and LN3D3 Fabs in a buffer containing 20 mM Tris-HCl pH8.0 and 150 mM NaCl on ice for 1 h. The complex was further purified on a Superdex 200 Increase 10/300 GL column (Cytiva) before cryo-EM grid preparation. Cryo-EM grids were prepared on a Thermo Scientific Vitrobot Mark IV at 4°C and 100% humidity. A total of 3.5 μL purified complex was applied to a freshly glow-discharged Cu 200 mesh R1.2/1.3 holey carbon grid (Quantifoil). After incubation for 20 seconds, the grids were blotted for 2 seconds at 100% humidity and 4 °C, and plunge-frozen in liquid ethane.

### Cryo-EM data collection and image processing

All data were collected using the CRYO ARM 300 electron microscope (JEOL, Japan) equipped with a K3 direct electron detector (Gatan, USA). Cryo-EM movies were recorded automatically using Serial-EM software in a super-resolution mode with a pixel size of 0.475 Å/pixel at a calibrated magnification of 50,000× over a defocus range of −0.5 to −2.5 µm. Data were collected at a frame rate of 40 frames per second with a total electron dose of 40 e/Å^2^.

Recorded movies were inputted into cryoSPARC for patch motion correction and CTF estimation ^50^. 4,784 micrographs were selected for further data processing. Particles were picked using the Topaz picker. 2,248,175 particles were extracted for 2D classification using a particle box size of 300 pixels. After two rounds of 2D classification, 1,207,684 particles were selected for two rounds of heterogeneous refinement. One class (145,635 particles) from the second round of heterogeneous refinement with good features was selected for nonuniform refinement (NU-refinement), yielding an overall resolution of 3.01 Å map.

### Model building

For the cryo-EM structure, the NiV_M_ G head domain model and AlphaFold2-predicted Fab models of variable regions were docked into the cryo-EM map using Chimera. For crystal structure, initial models were fitted into the density in Coot. Iterative model building and refinement were performed in Coot ^51^ and PHENIX ^52^. The data collection and refinement statistics for the final models are listed in Tables S1 and S2.

### Fusion inhibition assay

A dual-functional split-reporter system, which includes RL-DSP1-7 and RL-DSP8-11 expression vectors, was used for the fusion inhibition assay as described previously^33,43^. Briefly, 293T cells were seeded into 6-well plates one day before transfection. Effector cells were co-transfected with the full-length NiV_M_ G, F, and RL-DSP1-7 expression vectors, while the target cells were transfected with the RL-DSP8-11 expression vector. 6 h post-transfection, the effector and target cells were trypsinized and mixed into 96-well plates. NiV G mAbs/Fabs diluted in DMEM containing 10% FBS were added to the cells. Cells treated with DMEM containing 10% FBS without mAbs/Fabs were used as controls. For luciferase activity detection, after approximately 14-16 h of incubation, the cell culture medium was discarded and replaced with fresh DMEM containing 10% FBS and 20 μM EnduRen live-cell substrate (Promega, Cat#: E6482). The cells were incubated for at least 2 h, and live-cell luciferase activity induced by glycoprotein-mediated cell-cell fusion was detected using a Varioskan LUX (Thermo Scientific). For GFP detection, the cells were fixed by 4% PFA after 24-36 h of incubation, and the nuclei were stained with Hoechst 33342 (Thermo Scientific, R37165). Images of the same position in different experimental wells were captured using CTL-S6 Universal M2.

### Animal experiments-Syrian hamsters

Six-week-old female Syrian hamsters from Beijing Vital River Laboratories were randomly divided into five groups (n=6). For prophylactic treatment, hamsters were administered the indicated mAbs at 30 mg/kg via the intraperitoneal (i.p.) route, followed by a challenge with 1000 LD_50_ of NiV Malaysia strains via i.p. injection 24 h later. For therapeutic treatment, hamsters were challenged with 1000 LD_50_ NiV_M_ via i.p. injection. Then, hamsters were injected with tested NiV G mAb or isotype control mAb S2A5 (an anti-SFTSV mAb ^33^) twice at a dosage of 15 mg/kg through the i.p. route, once on day 1 and again on day 3 post-challenge. Hamsters were monitored daily post-challenge for survival over three weeks and for weight changes over two weeks.

## Data and code availability

The cryo-EM and crystal structures have been deposited to the electron microscopy data bank and protein data bank with accession numbers EMD-63395, PDB-9LUE (NiV G^H^-S1E2-S2B10-LN3D3 complex), and PDB-9LU3 (NiV G^H^-LN1F9 Fab complex). Other source data are available from the corresponding authors upon request.

## Acknowledgments

We are particularly grateful to the running team (Ge Gao, Yun Peng and Miaoyu Chen) of National Biosafety Laboratory, Wuhan, Chinese Academy of Sciences for their assistance. We thank the Center for Instrumental Analysis and Metrology of Wuhan Institute of Virology for providing technical assistance. We thank the staff of the BL02U1 and BL10U2 beamlines at Shanghai Synchrotron Radiation Facility for assistance during X-ray diffraction data collection. We appreciate the National Virus Resource Center for the important reagents. This work was supported by the National Key Research and Development Program (2022YFC2604100) and Wuhan Natural Science Foundation exploration program (202404080120225) to H.Z., the CAS Pioneer Hundred Talents Program to Z.D, and the Strategic Priority Research Program of the Chinese Academy of Sciences (XDB0490000 to S.C.).

## Author contributions

H.Z., Z.D. and S.C. conceived the project. D.Z. and R.C. conducted biochemical preparations and functional assays with the help of G.Z., and X.L.; Y.Y. and H.L. performed BSL-4 and ABSL-4 experiments. Y.W. and Z.D. determined the cryo-EM structure. W.K. determined the crystal structure. H.Z., D.Z., and Z.D. analyzed the data and wrote the manuscript with input from all authors.

